# Evaluating the efficacy of enzalutamide and the development of resistance in a preclinical mouse model of type-I endometrial carcinoma

**DOI:** 10.1101/2019.12.06.868182

**Authors:** Christopher S. Koivisto, Melodie Parrish, Santosh B. Bonala, Soo Ngoi, Adrian Torres, James Gallagher, Rebekah Sanchez-Hodge, Victor Zeinner, Georges J. Nahhas, Bei Liu, David E. Cohn, Floor J. Backes, Paul J. Goodfellow, Helen M. Chamberlin, Gustavo Leone

## Abstract

Androgen Receptor (AR) signaling is a critical driver of hormone-dependent prostate cancer and has also been proposed to have biological activity in female hormone-dependent cancers, including type I endometrial carcinoma (EMC). In this study, we evaluated the preclinical efficacy of a third-generation AR antagonist, enzalutamide, in a genetic mouse model of EMC, *Sprr2f-Cre;Pten^fl/fl^*. In this model, ablation of *Pten* in the uterine epithelium leads to localized and distant malignant disease as observed in human EMC. We hypothesized that administering enzalutamide through the diet would temporarily decrease the incidence of invasive and metastatic carcinoma, while prolonged administration would result in development of resistance and loss of efficacy. Short-term treatment with enzalutamide reduced overall tumor burden through increased apoptosis but failed to prevent progression of invasive and metastatic disease. These results suggest that AR signaling may have biphasic, oncogenic and tumor suppressive roles in EMC that are dependent on disease stage. Enzalutamide treatment increased Progesterone Receptor (PR) expression within both stromal and tumor cell compartments. Prolonged administration of enzalutamide decreased apoptosis, increased tumor burden and resulted in the clonal expansion of tumor cells expressing high levels of p53 protein, suggestive of acquired *Trp53* mutations. In conclusion, we show that enzalutamide induces apoptosis in EMC but has limited efficacy overall as a single agent. Induction of PR, a negative regulator of endometrial proliferation, suggests that adding progestin therapy to enzalutamide administration may further decrease tumor burden and result in a prolonged response.

## Introduction

The role of androgen receptor (AR) in tumorigenesis has been primarily studied in prostate cancer, however recent data suggest a role for AR signaling in other cancer types, particularly breast cancer, ovarian cancer, and endometrial cancer. Within endometrial cancer, the potential role of AR signaling is unclear. Knockout of AR in mice results in reduced uterus size and decreased responsiveness to the proliferative effects of estrogen[1]. Women with elevated circulating androgens have a 2-3 fold increased risk of developing endometrial cancer and administration of synthetic androgens to mice results in endometrial hyperplasia[2]. In humans, multiple studies observed increased AR protein expression in endometrial hyperplasia and cancer, with low-grade endometrioid, or type-I carcinomas more commonly positive than higher grade histologies[3–6]. Furthermore, positive AR expression has been associated with improved prognosis relative to AR-negative tumors[3,6]. Interestingly, increased AR expression was observed in metastatic tumors compared to matched primary tumors[3,4]. Clearly, the role of AR signaling in endometrial carcinogenesis is complex and may have different functions between early and late stages of cancer progression.

Loss of PTEN tumor suppressor function is an important driver of type I endometrial carcinoma in humans[7,8]. Ablation of *Pten* in the uterine tissue of mice is sufficient to drive endometrial carcinoma[9,10,11]. Specific deletion of *Pten* in endometrial glands of *Sprr2f-Cre;Pten^fl/fl^* mice models type I endometrial carcinoma[11]. By three months of age, 100% of *Sprr2f-Cre;Pten^fl/fl^* females have hyperplastic uterine remodeling and show malignant tumors confined within the endometrial compartment *(in-situ* carcinoma). Approximately 50% of mice develop endometrial tumors that invade into the myometrium (invasive carcinoma). A small but significant proportion of mice (~5%) have concurrent metastases, most commonly in the regional lymph nodes, but occasionally also within the ovarian parenchyma.

Enzalutamide is an inhibitor of AR signaling that prevents the nuclear translocation of Androgen Receptor and its sequence-specific binding to DNA, resulting in cellular apoptosis. *PTEN* loss is a frequent genetic alteration in prostatic cancer that results in activation of AKT signaling[12]. Furthermore, exogenous expression of phosphorylated AKT and AR in epithelial cells from the mouse prostate have a pronounced synergistic effect in promoting cell growth in both intact and castrated animals[13]. In mouse models of prostate cancer, oral administration of enzalutamide at doses of 1, 10 and 50 mg/kg/day for 4-12 weeks results in a reduction of tumor growth and disease progression[14]. Enzalutamide is now approved for use in castration-resistant prostate cancer[15]. To evaluate the role of AR signaling in the pathogenesis of type I endometrial carcinoma and the potential efficacy of enzalutamide in treating uterine cancer, we evaluated the effects of enzalutamide on tumor progression in *Sprr2f-Cre;Pten^fl/fl^* mice. We entertained the hypothesis that impairing AR signaling through enzalutamide treatment would decrease tumor burden and progression of invasive and metastatic disease. Here, we demonstrate that short-term enzalutamide, as a single-agent, results in increased apoptosis and a dose-dependent decrease in tumor burden. Prolonged enzalutamide treatment, however, failed to sustain tumor reduction and was associated with decreased apoptosis and clonal expansion of cells with overexpression of p53 protein, indicating eventual resistance to treatment.

## Materials and methods

### Mouse models and study design

Sprr2f-Cre mice were purchased from the NCI repository (Strain No. 01XNA)[16]. Pten-floxed mice were generated using standard homologous recombination cloning techniques as previously reported[17]. *Sprr2f-Cre;Pten^fl/fl^* mice were generated by crossing *Sprr2f-Cre;Pten^fl/+^* males and females. All mice were in a pure FVB background and only females were used for this study. Genotyping of offspring was performed on tail DNA by standard PCR using allele-specific primers listed in table 1. Mouse usage and experiments were approved by the Institutional Animal Care and Use Committee at The Ohio State University. Mice were housed, five or less animals per cage, under normal husbandry conditions in a vivarium with a 12-hour light/dark cycle. Female *Sprr2f-Cre;Pten^fl/fl^* mice were weaned at 21 days and maintained on standard chow until 6 weeks of age to allow for uterine tumors to grow. At 6 weeks of age, mice were randomly assigned to one of three treatment groups (control, low-dose enzalutamide, or high-dose enzalutamide) and fed an exclusive diet *ad lib* containing vehicle or drug for 8- or 16-weeks. Food consumption and body weights of mice were recorded twice weekly while on treatment. At the end of treatment, mice were euthanized by asphyxiation with carbon dioxide and dissected. The female reproductive tract was isolated. The uterus was separated from the ovaries/oviducts and cervix/vagina and weighed. Spleen and thymus were also weighed. Three-quarters of the spleen was saved for isolation of splenocytes. Pieces of liver were flash frozen in liquid nitrogen and stored at −80C. The remaining tissue was preserved in 10% neutral buffered formalin (Fisher Scientific; 23-245-685) for 24-48 hours and rinsed in 70% ethanol. Genotyping of study mice was confirmed using liver DNA.

**Table 1.**
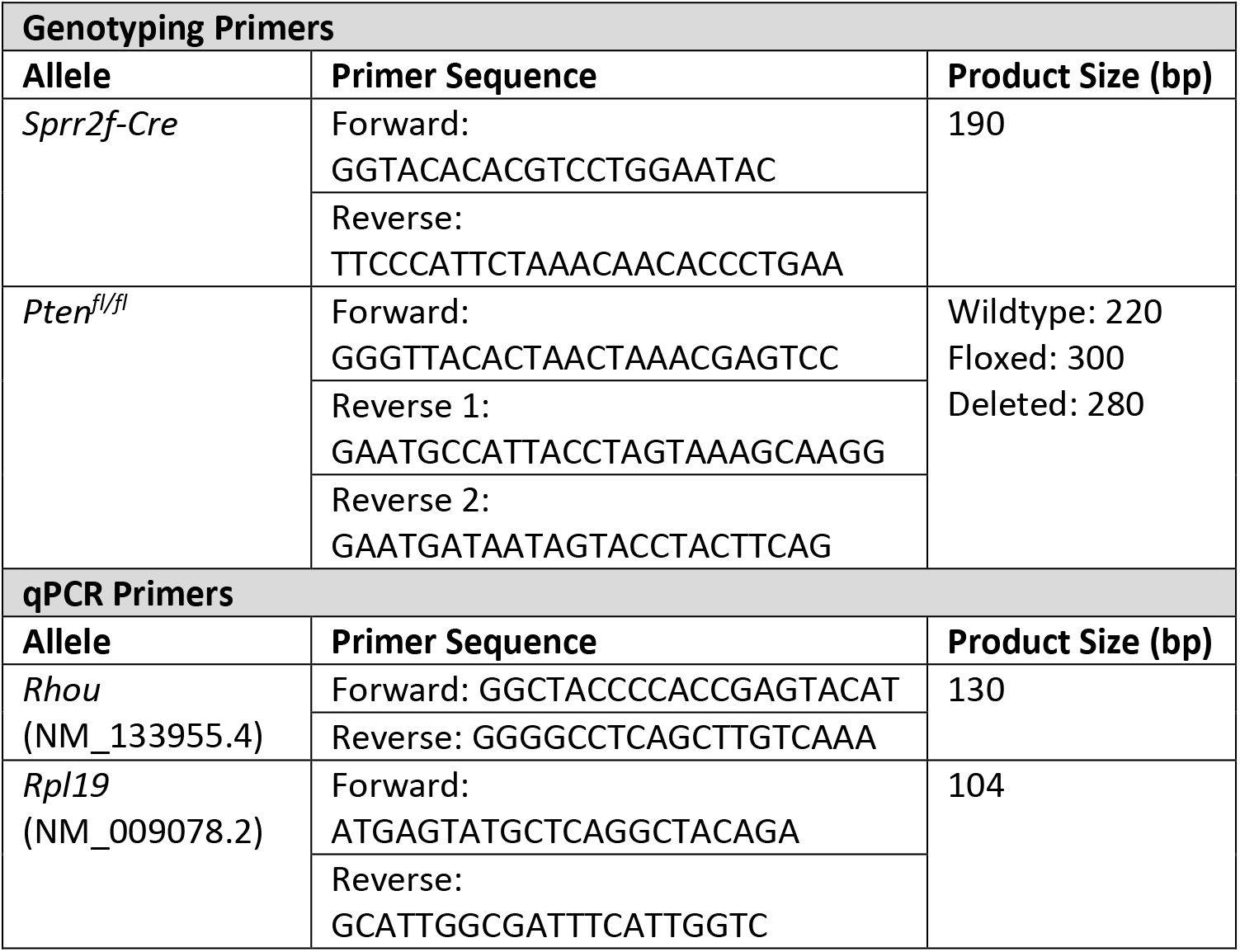
PCR Primers

### Diet formulation

Diets containing enzalutamide were specially formulated by Research Diets, Inc. (New Brunswick, NJ). Enzalutamide was shipped directly to Research Diets by the study sponsor. Low-dose and high-dose enzalutamide diets were generated by respectively adding 171mg or 429mg of enzalutamide per kg of vehicle diet AIN-76A (D10001). These formulations were estimated to deliver 0, 20 or 50mg/kg of drug per day to study mice. Diets were stored at 4C in the dark until fed and residual food was changed out weekly.

### Tissue preparation for histology and pathology analysis

Uterus, vagina, ovaries, liver, lung, sublumbar lymph nodes, perirenal lymph nodes, spleen and thymus from all study mice were fixed in 10% neutral buffered formalin for 24-48 hours at room temperature, then transferred to 70% ethanol. Tissue was processed for routine histology in an automatic processor, embedded in paraffin, cut into 4μm sections and stained with hematoxylin and eosin (H&E). Uterine tissue was examined microscopically to document the presence of *in situ* carcinoma or invasive carcinoma. *In situ* carcinoma is characterized by the expansion of endometrial glands combined with loss of cellular polarity and cribiform growth patterns. *In situ* carcinomas expand and replace the existing endometrial stroma but remained confined to the boundaries of the endometrial layer. In contrast, invasive carcinomas exhibit similar cellular morphologies but extend beyond the confines of the endometrium and invade into underlying myometrium and vasculature. Lymph nodes, ovaries, liver and lung were also examined microscopically for the presence of metastases.

### Immunostaining and quantification

Four μm thick, unstained sections were placed on positively charged slides. Immunostaining was performed on a Bond Rx (Leica) or Ventana Discovery Ultra (Roche) autostainer. Protocols, primary antibodies and dilutions are summarized in table 2. Immunostained slides were scanned using Vectra Polaris (Perkin Elmer)instrumentation at 20x resolution. Three 2×2 or 3×3 fields, representing the highest expression levels for each sample, were manually selected for multispectral image acquisition. Quantification of immunohistochemistry was performed on multispectral images using Inform software (Perkin Elmer). Epithelial and stromal compartments were manually annotated and analyzed independently. Quantification of Ki67 and CC3 expression were determined by the percent of positive nuclei within a given compartment. Quantification of ER, PR and GR expression was performed by measuring the nuclear staining intensity. The spectrum of signal intensity was divided into 10 bins, from lowest (bin 0) to highest (bin 9). Each nucleus was assigned to a bin and a modified H-score was calculated for each image. H-scores were calculated as shown in the formula below resulting in a value between 0-900, where zero is absolute negative expression and 900 is all cells expressing the highest level of intensity.

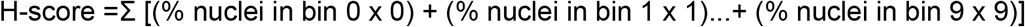

**Table 2.** Antibodies for immunohistochemistry

| Antigen Target | Manufacturer | Catalog # | Concentration | Antigen Retrieval Method |
| --- | --- | --- | --- | --- |
| Ki67 | Abcam | AB1667 | 1:400 | EDTA |
| Cleaved Caspase 3 | Cell Signaling | 9579 | 1:500 | EDTA |
| Progesterone Receptor | Thermo | RM-9102-SO | 1:20 | CC1 (Tris-EDTA) |
| Estrogen Receptor *alpha* | Abcam | AB32063 | 1:2000 | Citrate |
| p53 | Leica | PA0057 |  | EDTA |
| Glucocorticoid Receptor | Cell Signaling | 12401 | 1:400 | Citrate |

### Isolation of splenocytes and flow cytometry

Splenocytes were prepared from the spleens of study mice using a previously described protocol with the following modifications[18]. Spleens were placed onto a 70μm cell strainer with 500μl of red blood cell (RBC) lysis buffer, then crushed through the strainer and into a 50ml tube using a rubber syringe plunger. Additional RBC lysis buffer was added as necessary to get all tissue through the strainer. Cells were incubated in RBC lysis buffer for 3 minutes at room temperature. Following RBC lysis, 10ml of fluorescent activated cell sorting (FACS) buffer was added to tubes to wash cells followed by centrifugation to pellet cells. Cells were washed twice then resuspended in 2ml of freezing media (DMEM media with 20% fetal bovine serum + 10% dimethylsulfoxide) and stored at −80C in 1ml aliquots.

For flow cytometry, frozen splenocytes were thawed and diluted in 10ml of FACS buffer and filtered through 20μm nylon mesh filters to remove debris. Cells were pelleted, resuspended in 50ul of 1x phosphate buffered saline containing a cocktail of primary antibodies (Table 3) and incubated on ice in the dark for 30 minutes. Cells were washed and resuspended in FACS buffer for flow cytometry.

**Table 3.**
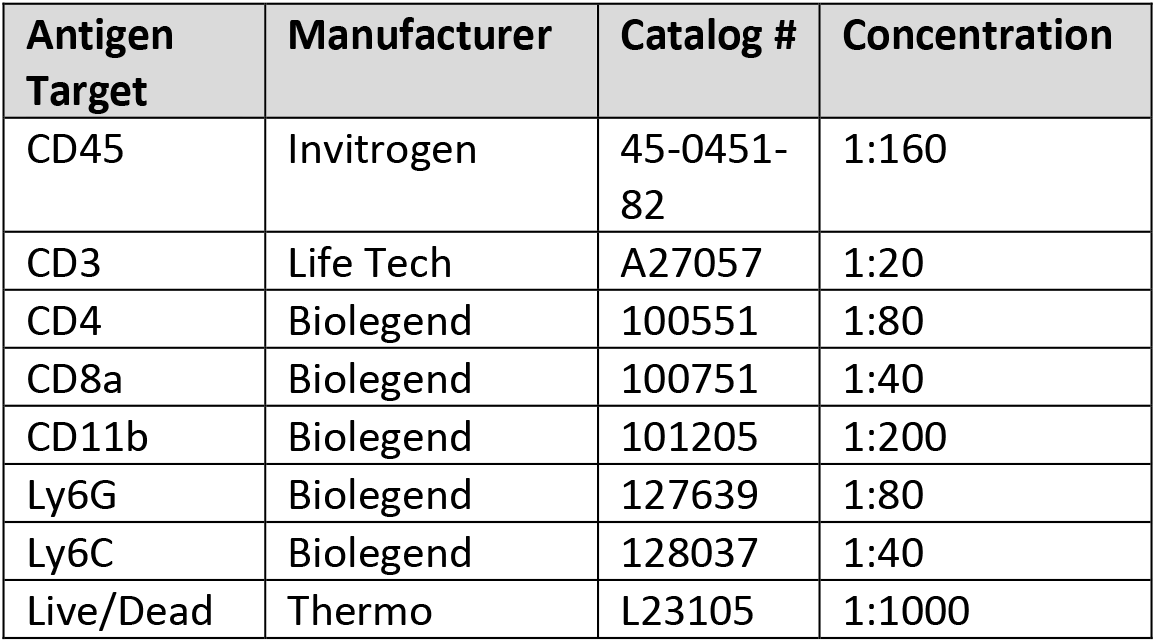
Antibodies for flow cytometry

| Antigen Target | Manufacturer | Catalog # | Concentration |
| --- | --- | --- | --- |
| CD45 | Invitrogen | 45-0451-82 | 1:160 |
| CD3 | Life Tech | A27057 | 1:20 |
| CD4 | Biolegend | 100551 | 1:80 |
| CD8a | Biolegend | 100751 | 1:40 |
| CD11b | Biolegend | 101205 | 1:200 |
| Ly6G | Biolegend | 127639 | 1:80 |
| Ly6C | Biolegend | 128037 | 1:40 |
| Live/Dead | Thermo | L23105 | 1:1000 |

### Statistical Analyses

We employed a mixed model for repeated measures to compare mean body weight gains and mean food consumption between the three dose groups for each timepoint. This approach allows for analysis of unbalanced data where a different number of measurements were taken from different animals at any given time. Mean uterus:body weight ratios for each dose group were analyzed by one-way ANOVA for each stage of the estrous cycle and timepoint, with post-hoc Bonferonni tests to determine which individual dose groups were different. The incidence of progressive (invasive or metastatic) versus nonprogressive (*in situ*) EMC was compared between the three dose groups at each timepoint using Chi-square analysis. For gene expression, mean delta-CT values from control and high-dose groups at the 8-week timepoint were compared by Student’s T-test. For immunohistochemistry, mean H-scores from vehicle control and high-dose groups were compared by Student’s T-test for each timepoint. For flow cytometry, the mean percentage of gated cells expressing a designated marker was compared between vehicle and high-dose treatment groups by Student’s T-test for the 8-week timepoint. An alpha =0.05 was used for all statistical tests.

## Results

### Administration of enzalutamide has no effect on food consumption or body weight in Sprr2f-Cre; Pten^fl/fl^ mice

To test the efficacy of enzalutamide as a novel therapy for uterine carcinoma, we utilized female *Sprr2f-Cre; Pten^fl/fl^* mice, a model of type I endometrial carcinoma that were previously shown to develop endometrial tumors by 6 weeks of age[19]. The experimental design is summarized in Figure 1A. Six-week-old *Sprr2f-Cre; Pten^fl/fl^* female mice were subjected to a customized diet containing either control, low-dose or high-dose enzalutamide. Mice were group housed and fed treatment diets *ad libitum* for either eight or sixteen weeks. Food consumption by cage and body weights of individual mice were measured approximately every three days. There were no adverse differences in food consumption or body weight across the three treatment groups, suggesting that enzalutamide in the diet had no effect on food palatability or toxicity (Figure S1A and S1B). However, we did observe that mice fed high-dose enzalutamide had mildly increased body weights compared to control mice at both timepoints, possibly indicating that AR inhibition in females may increase weight gain. Quantitative polymerase chain reaction (qPCR) evaluation of *Rhou* expression, a classical AR-target gene, in the liver resulted in decreased AR-dependent gene expression (Figure S1C), confirming enzalutamide had a systemic effect on Androgen Receptor (AR) function.

**Figure 1.**
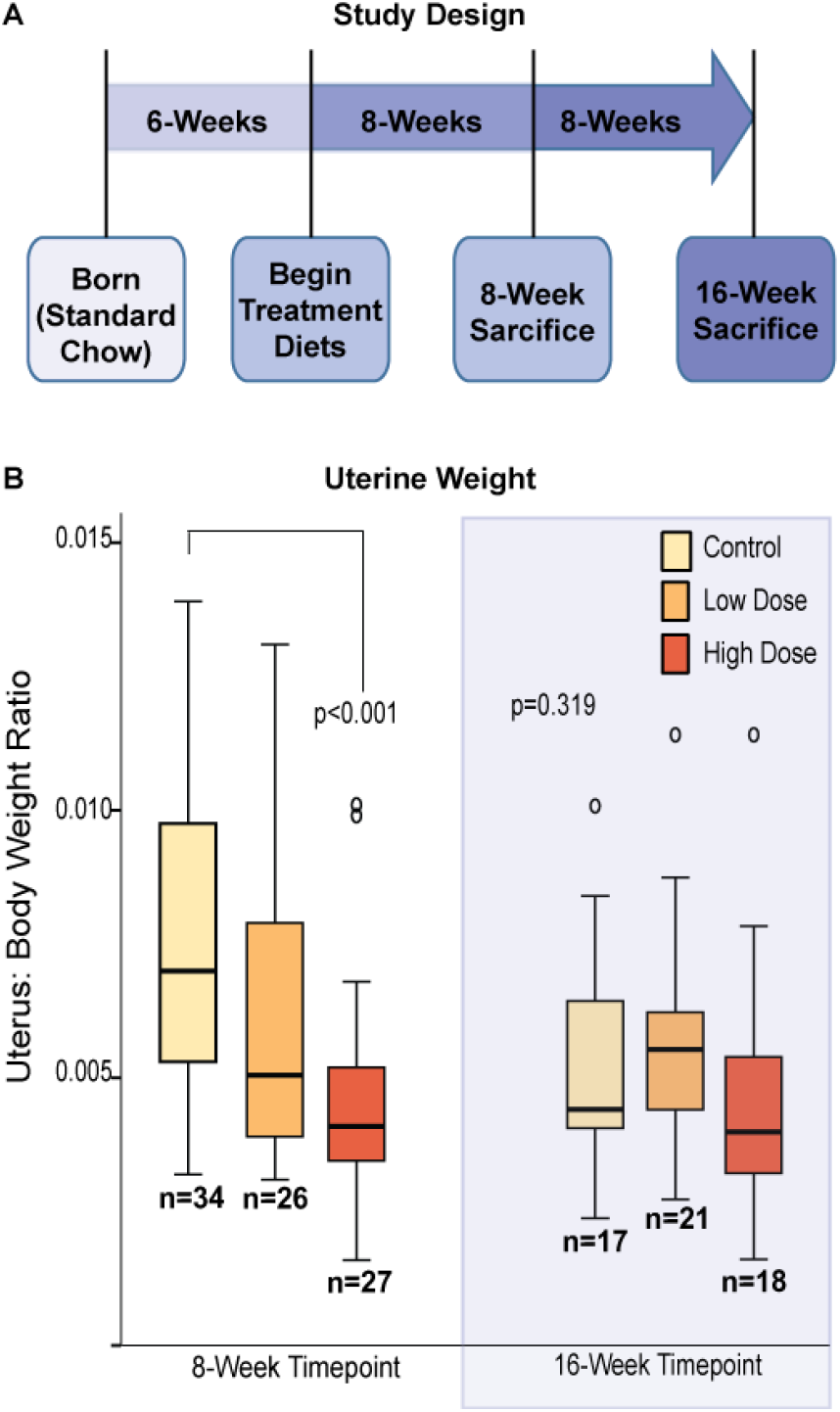
Effects of enzalutamide on uterine weights at 8- and 16-week timepoints. (A) Summary of study design. (B) Boxplots of individual uterine:body weight ratios. Circles represent outliers that are >1.5 box-lengths beyond the median. Statistical analysis comparing mean ratios for each treatment group and each timepoint performed by one-way ANOVA with post-hoc Bonferroni testing.

### Short-term enzalutamide administration increases tumor cell apoptosis and reduces endometrial tumor burden in Sprr2f-Cre;Pten^fl/fl^ mice

Within the *Sprr2f-Cre;Pten^fl/fl^* model, nearly all endometrial glands are deleted for *Pten*. Not surprisingly, malignant transformation appears as a polyclonal event affecting the majority of glands[11,20]. In this model, we assessed uterine weights as a surrogate of endometrial tumor burden following 8- or 16-weeks of enzalutamide therapy (Figure 1B). At the 8-week timepoint, we observed a dose-dependent decrease in uterine weights. However, by 16-weeks of treatment, the therapeutic benefits were lost. In conclusion, these data suggest that enzalutamide administration initially reduces tumor burden, while prolonged exposure leads to development of resistance to therapy.

Under normal physiological conditions, uterine weights change with each of the four stages of the estrous cycle in response to altering levels of circulating estrogen and progesterone. Using vaginal histology, we categorized individual mice into their respective stage of estrous (Supplemental Figure 1D-E) and compared uterine weights across each stage. Inadvertent cre-mediated deletion of *Pten* in the vaginal epithelium perturbed vaginal morphological features used for staging precluded analysis in a subset of female mice. As a result, there were an insufficient number of mice to be assessed at diestrus. This analysis showed that at 8-weeks, the dose-dependent reduction in unstaged uterine weights were reflected in females that were at estrus and metestrus (Supplemental Figure 1E). In contrast, enzalutamide had no significant impact on uterine weights at 16-weeks at any stage of the estrous cycle.

Unexpectedly, mice receiving low-dose enzalutamide had significantly higher uterine weights compared to control and high-dose groups (Supplemental Figure 1E) at proestrus. Proestrus is the proliferative phase of the estrous cycle for the endometrium and characterized by high levels of circulating estrogen and low levels of circulating progesterone. These results suggest that under-dosing enzalutamide during high estrogen signaling may result in undesired synergy for estrogen-dependent proliferation.

Studies of prostate cancer have attributed the therapeutic effects of enzalutamide to reducing tumor cell proliferation and increasing apoptosis[14]. Normal endometrium undergoes repetitive cycles of proliferation, differentiation and involution, which are all regulated by the sex hormones estrogen and progesterone. These same hormones can affect endometrial tumor cells as well. Using immunohistochemistry for Ki67 and cleaved Caspase 3 (CC3), we evaluated the effects of high-dose enzalutamide on proliferation and apoptosis, respectively, of malignant epithelium in *Sprr2f-Cre;Pten^fl/fl^* mice. To minimize the confounding effects of estrous cycle-dependent variation in proliferation and apoptosis, analysis was confined to *Sprr2f-Cre;Pten^fl/fl^* mice identified in proestrus. During proestrus, the proliferative phase of the estrous cycle, tumor cell Ki67 expression was similarly high in both control and high-dose treated mice at both timepoints (Figure 2). These results indicate that tumor cells, regardless of enzalutamide treatment, remain responsive to mitogenic hormone signaling associated with the estrous cycle. CC3 expression, on the other hand, was markedly increased in high-dose enzalutamide treatment at the 8-week timepoint relative to controls (Figure 3). Interestingly, the induction of apoptosis by enzalutamide dropped precipitously to control levels at the 16-week timepoint. These results indicate that short-term administration of enzalutamide reduced overall tumor burden through increasing apoptosis, without impacting proliferation. Resistance to long-term treatment with enzalutamide appears to be associated with a failure of tumor cells to undergo apoptosis.

**Figure 2.**
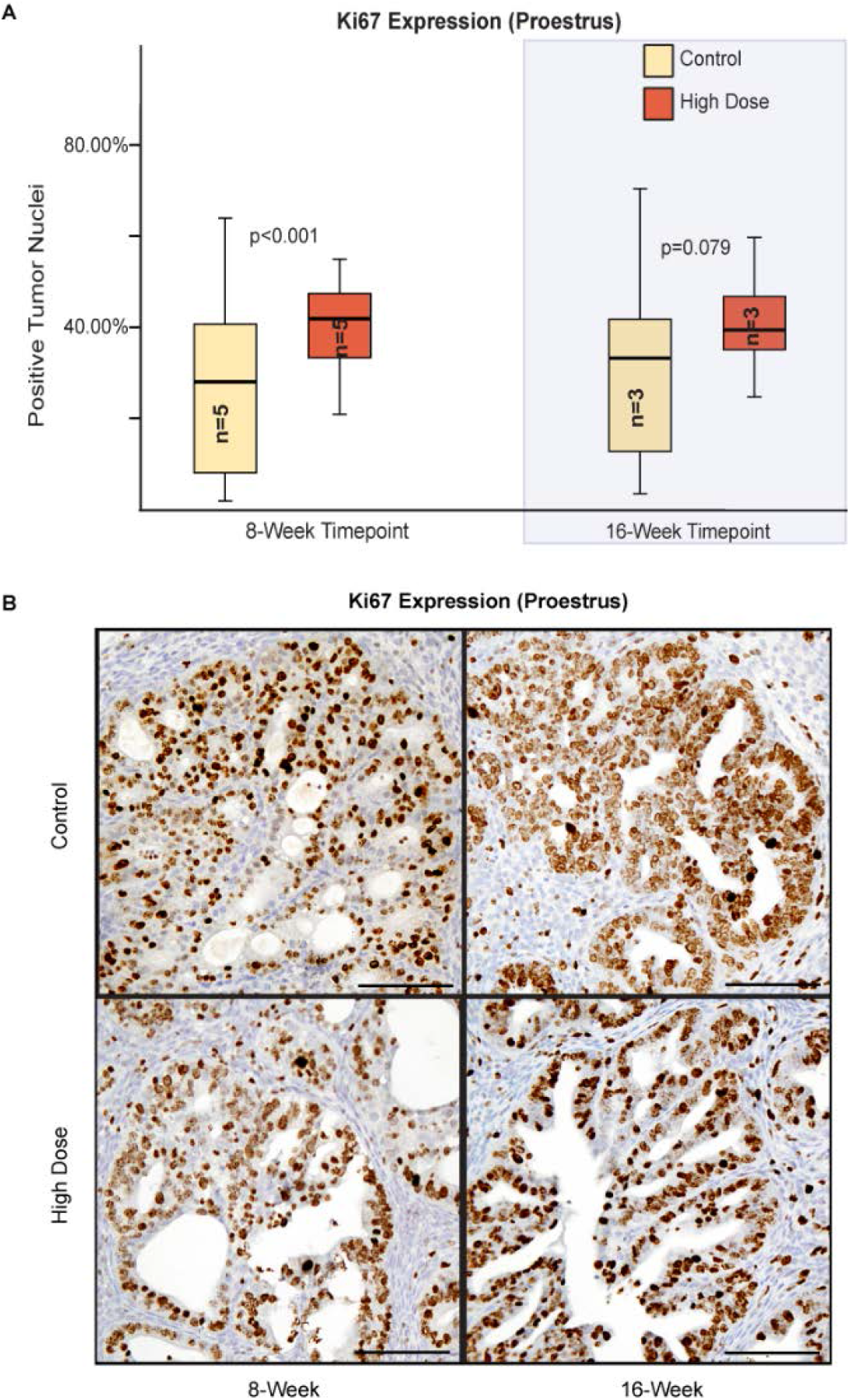
Ki67 expression by immunohistochemistry. (A) Boxplots quantifying mean percentage of positively stained cells expressing Ki67.Three 2×2 or 3×3 200x fields were selected from each mouse and quantified by digital image analysis. Only neoplastic glands were analyzed. Statistical analysis by Student’s T-test comparing mean percentage of positively stained tumor cells between vehicle control and high-dose groups at each timepoint performed. (B) Representative photomicrographs of Ki67 (brown) staining counterstained with hematoxylin. Scale bars: 100μm.

**Figure 3.**
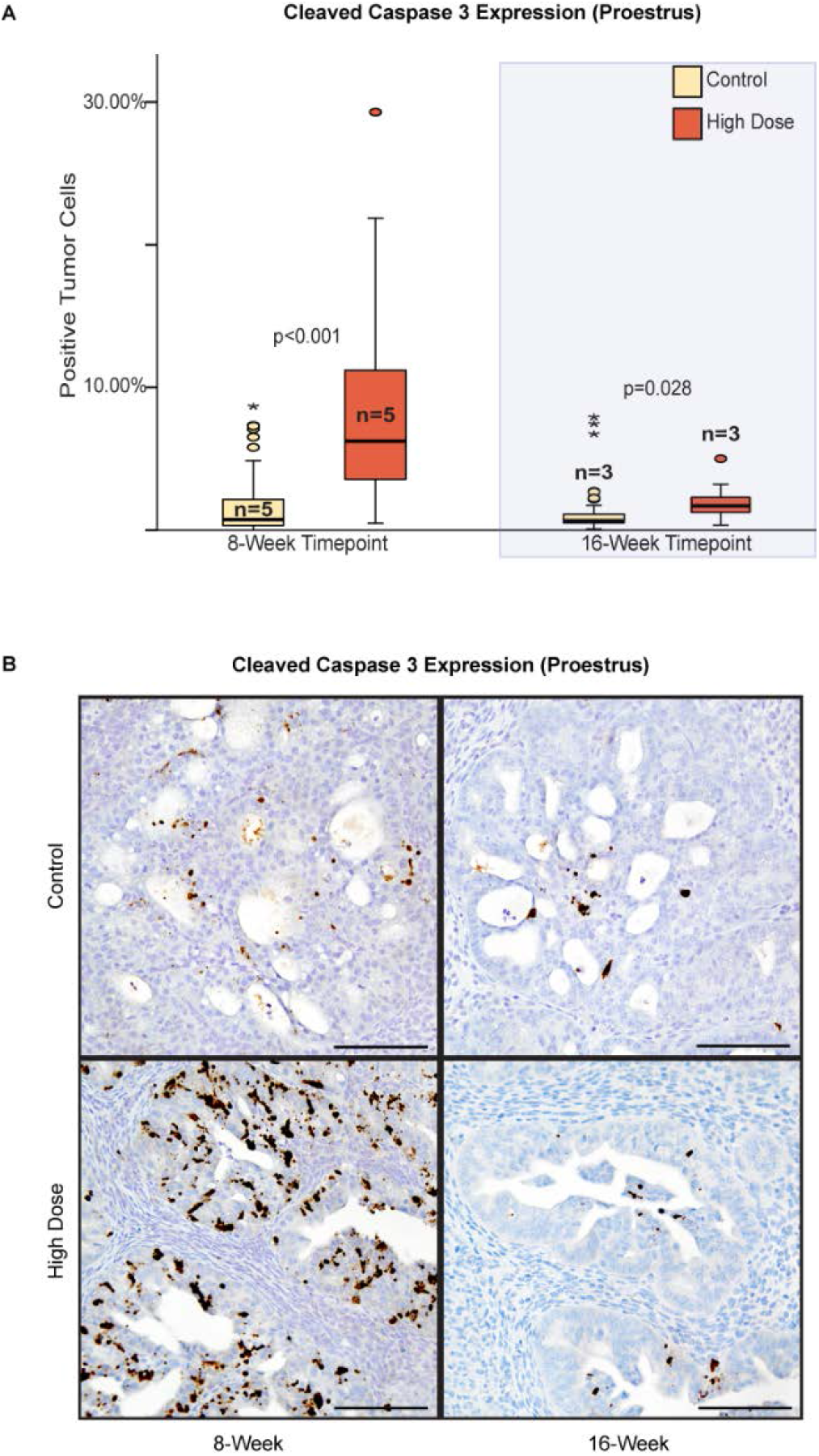
Cleaved Caspase 3 expression by immunohistochemistry. (A) Boxplots quantifying mean percentage of positively stained cells expressing CC3.Three 2×2 or 3×3 200x fields were selected from each mouse and quantified by digital image analysis. Only neoplastic glands were analyzed. Statistical analysis by Student’s T-test comparing mean percentage of positively stained tumor cells between vehicle control and high-dose groups at each timepoint. (B) Representative photomicrographs of CC3 (brown) staining counterstained with hematoxylin. Scale bars: 100μm.

### Enzalutamide administration is ineffective at preventing endometrial tumor progression in Sprr2f-Cre;Pten^fl/fl^ mice

Next, we characterized endometrial tumor morphology and documented the incidence of myoinvasion and metastasis, which are key features of cancer progression within the *Sprr2f-Cre;Pten^fl/fl^* mouse model[21]. Ovaries, sublumbar and perirenal lymph nodes were examined from all mice for incidence of metastatic carcinoma. *Sprr2f-Cre;Pten^fl/fl^* mice develop *in situ* carcinoma with 100% penetrance at an early age. *In situ* carcinomas are characterized by neoplastic expansion of the glandular compartment without invasion into the underlying myometrium. Tumor cells are well-differentiated with loss of polarity but minimal nuclear atypia. The incidence of *in situ* carcinoma, invasive carcinoma and metastatic carcinoma are summarized in Figure 4A. Mice had similar tumor morphology regardless of enzalutamide treatment (Figure 4B). At 8-weeks post treatment, there was an unexpected dose-dependent increase in the incidence of invasive and metastatic carcinoma (Figure 4A), even though enzalutamide led to a clear reduction in primary tumor burden (Figure 1B). Because of this discrepancy, we postulate that enzalutamide administration, and thus AR signaling, may have distinct and opposing functions during early and late stages of endometrial tumor progression.

**Figure 4.**
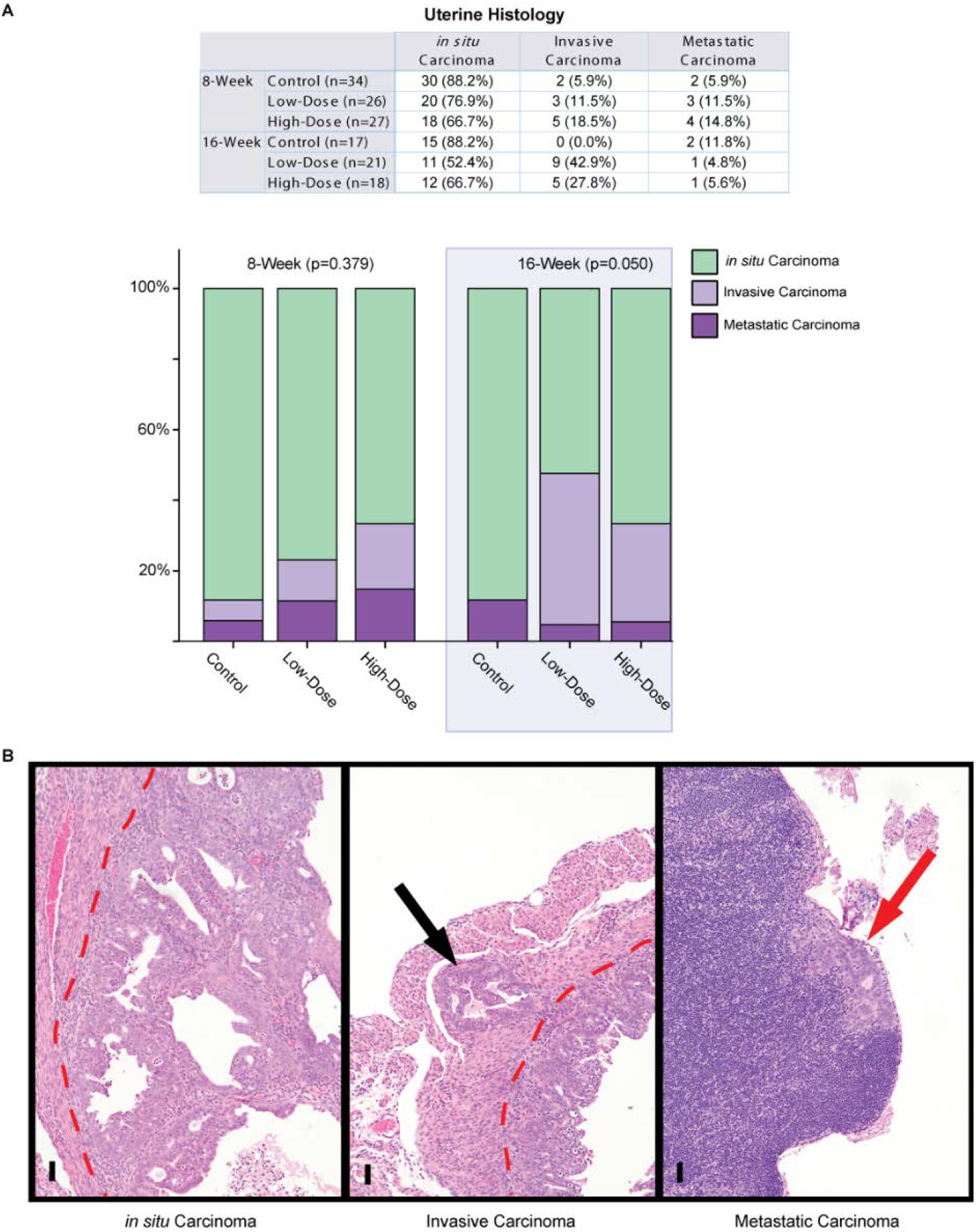
Incidence of uterine *in situ* carcinoma, invasive carcinoma and metastatic carcinoma. (A) Table and bargraphs summarizing tumor incidences. Statistical analysis by Chi-square comparing incidence of progressive (invasive or metastatic carcinoma) disease across the three dose groups for each timepoint. (B) Photomicrographs of representative tumor morphologies. Hematoxylin and Eosin. Red dashed line separates myometrium (left) from endometrium (right). Black arrow indicates myoinvasion of neoplastic cells. Red arrow indicates metastatic endometrial carcinoma within the subcapsular sinus of a lymph node. Scale bars: 50μm.

### Enzalutamide administration increased progesterone receptor expression within epithelial and stromal compartments

Endometrial growth is regulated through complex stromal-epithelial interactions that are mediated by circulating sex hormones estrogen and progesterone, which bind to their respective hormone receptors expressed in epithelial and stromal cells. We used Progesterone Receptor (PR) and Estrogen Receptor *alpha* (ER) specific antibodies, to evaluate the effects of high-dose enzalutamide on uterine tissue (epithelial and stromal) from mice in proestrus. Immunohistochemical analysis of 8-week treated mice showed that ER expression was unchanged in either tissue compartment relative to controls. In contrast, there was a moderate reduction in stromal ER expression following 16-weeks of enzalutamide treatment (Figure 5). The delayed reduction in stromal ER expression may be an indicator of enzalutamide resistance since ER expression in controls remained unaltered at the 8- and 16-week timepoints.

**Figure 5.**
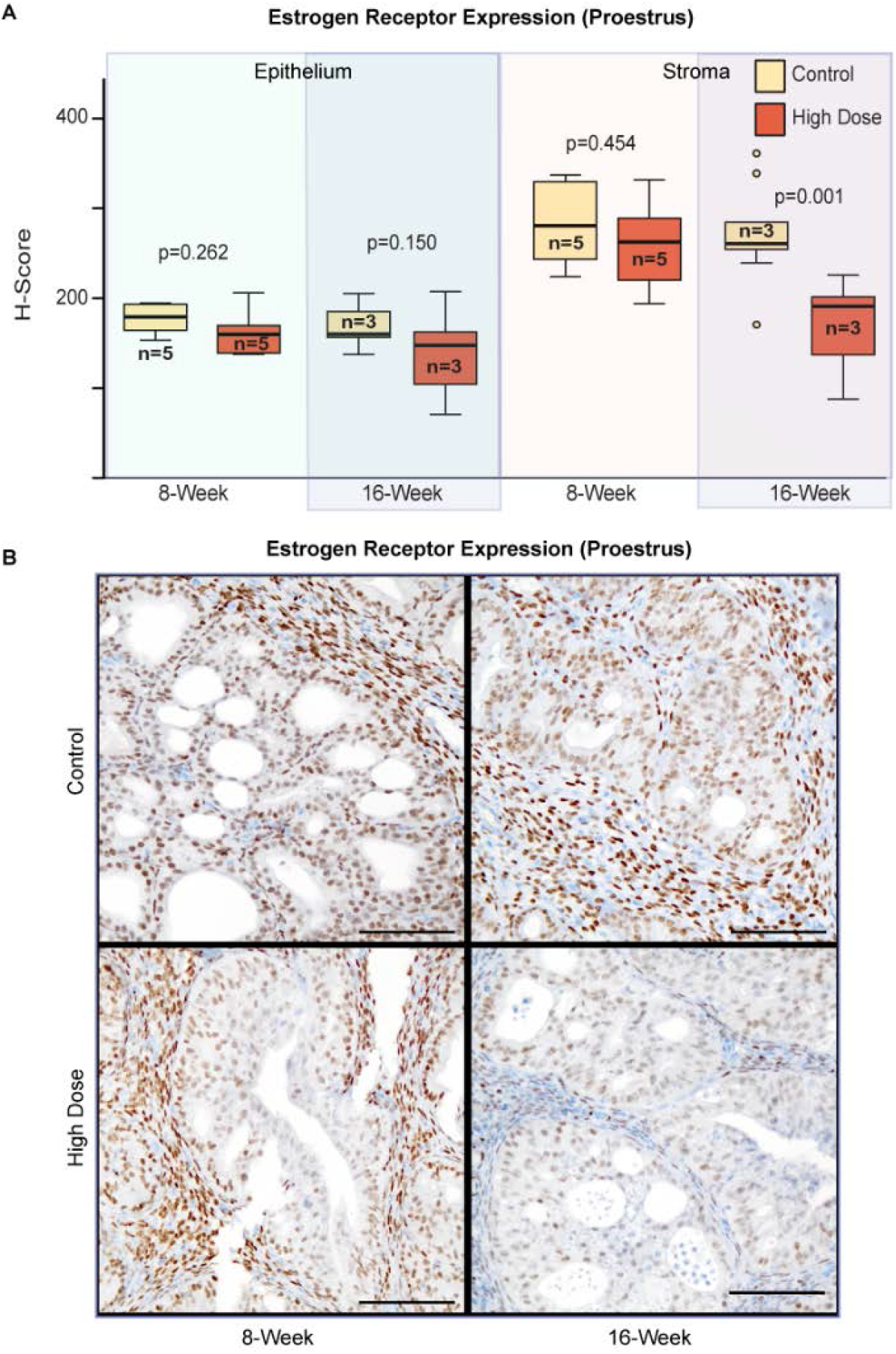
Expression of Estrogen Receptor *alpha* by immunohistochemistry. (A) Boxplots quantifying mean H-score of ER staining. Three 2×2 or 3×3 200x fields were selected from each mouse and quantified by digital image analysis. Statistical analysis by Student’s T-test comparing mean H-score between vehicle control and high-dose groups for each tissue compartment and each timepoint. (B) Representative photomicrographs of ER (brown) staining counterstained with hematoxylin. Scale bars: 100μm

Epithelial PR expression moderately increased at 8-weeks and remained elevated at 16-weeks post treatment (Figure 6). Stromal PR expression was unaltered at 8-weeks but markedly increased at 16-weeks. These results indicate that enzalutamide upregulates PR expression within the endometrial epithelium, and suggests that enzalutamide resistance may be associated with delayed upregulation of stromal PR.

**Figure 6.**
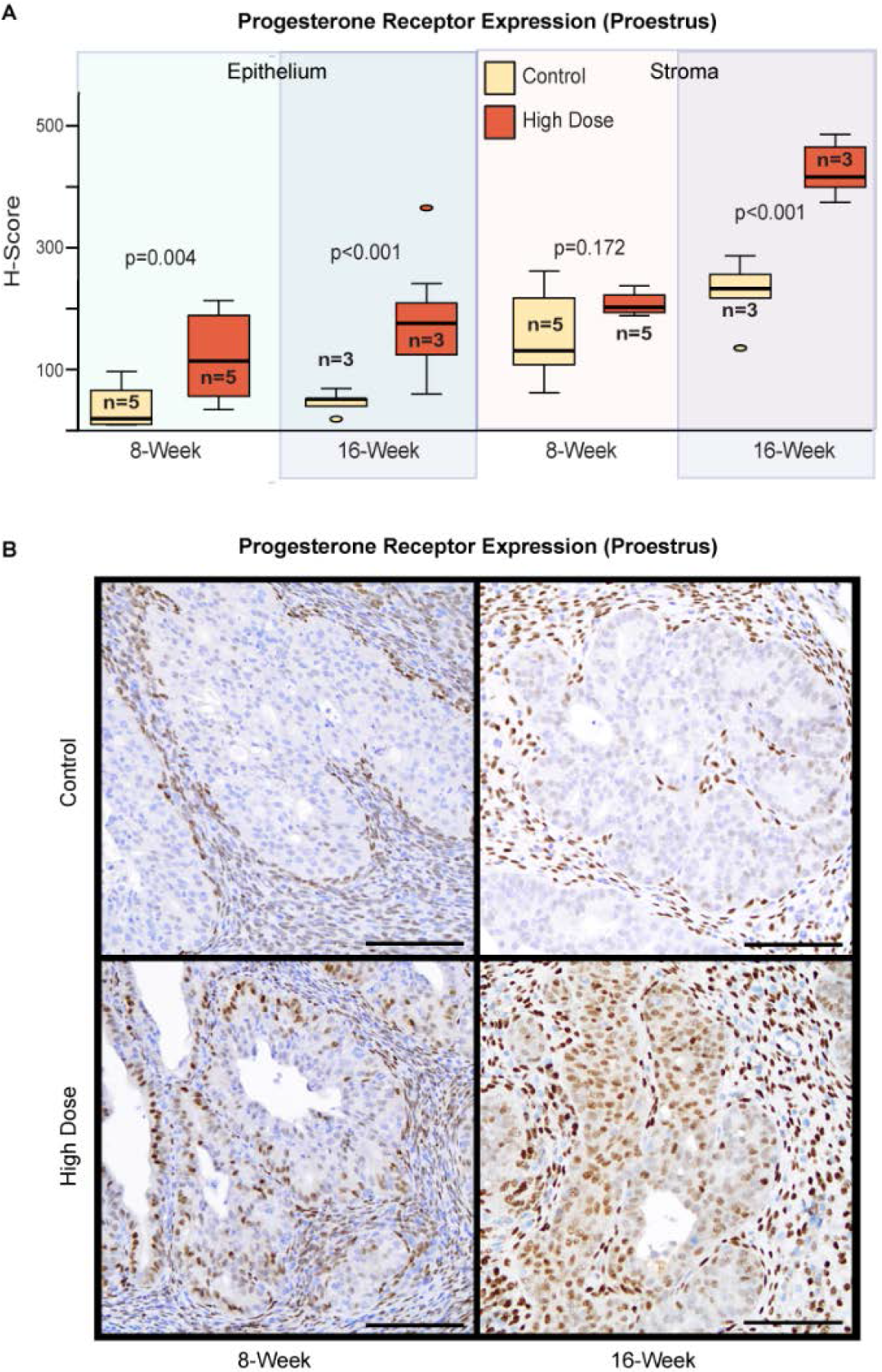
Expression of Progesterone Receptor by immunohistochemistry. (A) Boxplots quantifying mean H-score of PR staining. Three 2×2 or 3×3 200x fields were selected from each mouse and quantified by digital image analysis. Statistical analysis by Student’s T-test comparing mean H-score between vehicle control and high-dose groups for each tissue compartment and each timepoint. (B) Representative photomicrographs of PR (brown) staining counterstained with hematoxylin. Scale bars: 100μm

### Enzalutamide resistance in Sprr2f-Cre;Pten^fl/fl^ mice is associated with clonal upregulation of p53 protein

Previous studies on prostate cancer have proposed upregulation of Glucocorticoid Receptor (GR) as a marker of enzalutamide resistance[22]. We evaluated GR expression by immunohistochemistry in control and high-dose treated mice at 8- and 16-week timepoints. Enzalutamide induced GR expression within epithelial and stromal compartments at the 8-week timepoint (Figure 8). However, GR expression returned to control levels at the 16-week timepoint. These results suggest that the role of GR in enzalutamide resistance differs between endometrial cancer and prostate cancer.

Given the observed decrease in tumor cell death following long-term enzalutamide treatment, we hypothesized that long-term administration of enzalutamide might alter the p53 pathway to bypass apoptotic engagement and suppress tumor growth. Evaluation of p53 expression by immunohistochemistry showed that one-third of mice treated with high-dose enzalutamide for 16-weeks exhibited foci of clustered cells with uniformly intense p53 immunoreactivity (Figure 7). We termed this specific pattern of expression as “clonal”, which was distinct from the “random” pattern of p53 expression observed sporadically throughout the uteri. Clonal p53 expression was observed only in mice treated with high-dose enzalutamide for 16-weeks, whereas random p53 expression was observed across all groups. These results suggest that clonal overexpression of p53, likely indicative of acquired p53 mutations, may represent one mechanism for the observed enzalutamide resistance.

**Figure 7.**
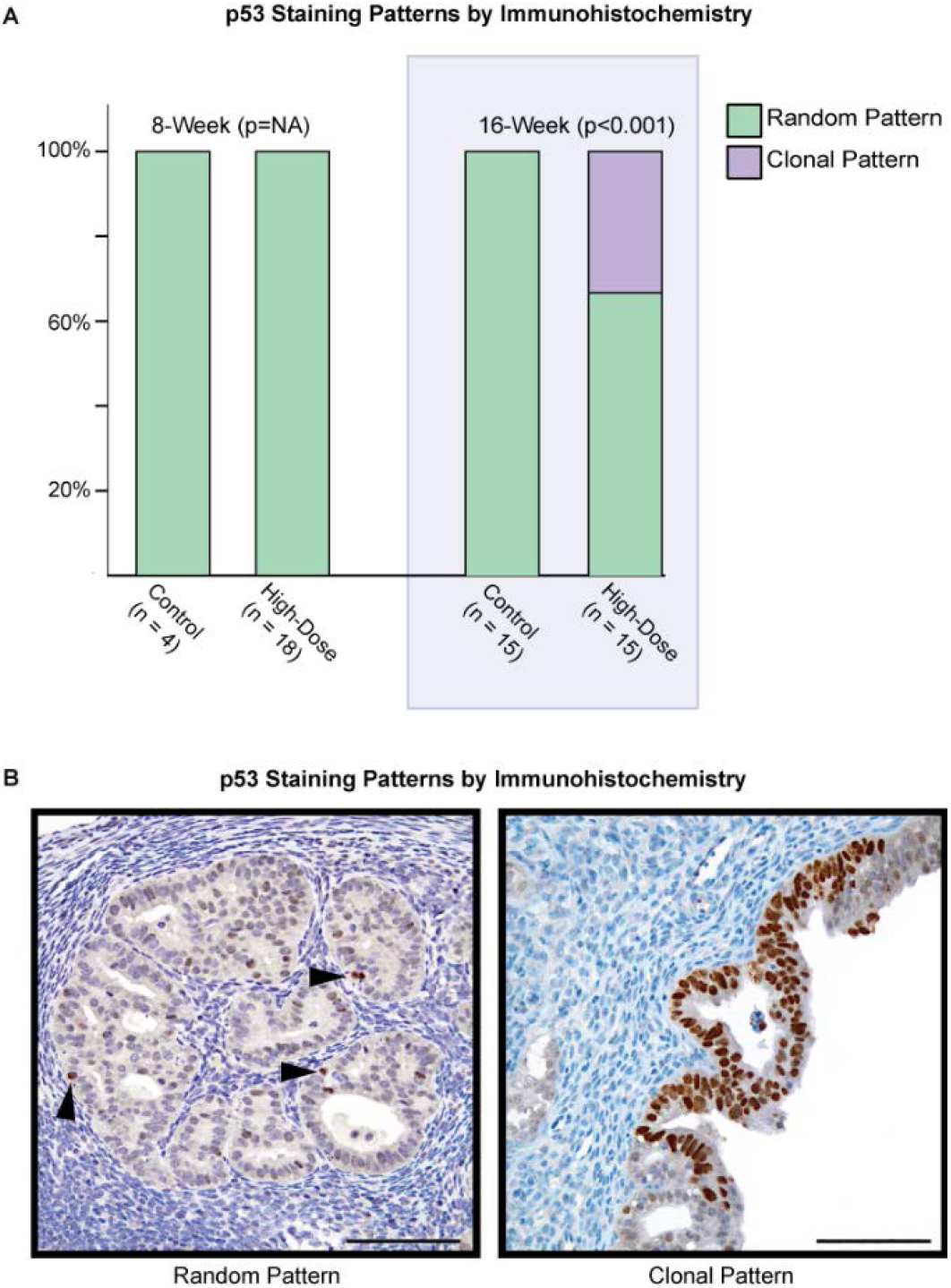
Expression of p53 by immunohistochemistry. (A) Barplots summarizing the p53 staining patterns observed in mouse uterine tumors. Statistical analysis by Chi-square comparing the incidence of clonal staining between vehicle control and high-dose groups for 16-week timepoint. (B) Representative photomicrographs of p53 (brown) staining counterstained with hematoxylin. Arrowheads point to random tumor cells expressing nuclear p53. Scale bars: 100μm

**Figure 8.**
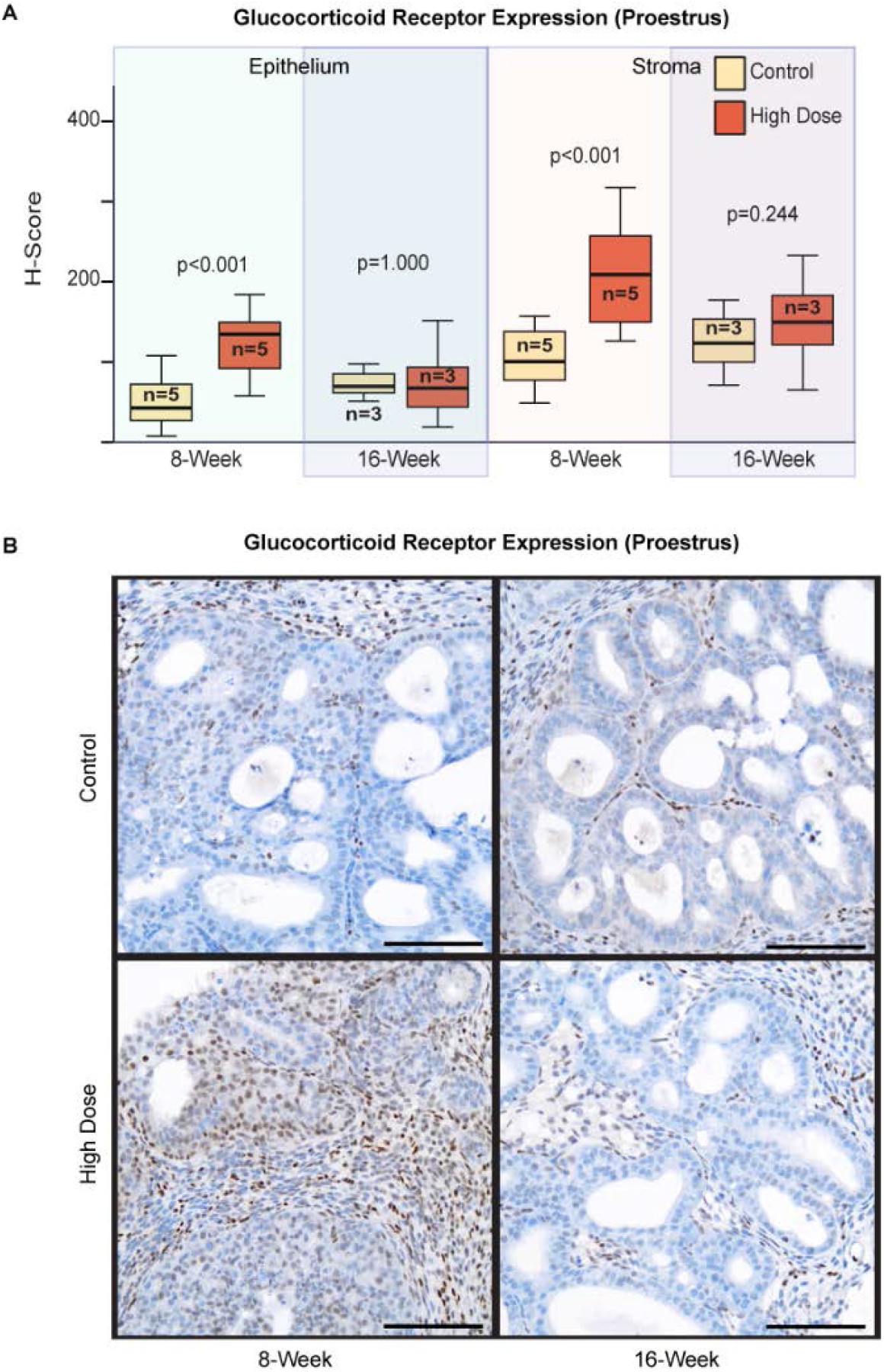
Expression of Glucocorticoid Receptor by immunohistochemistry. (A) Boxplots quantifying mean H-score of GR staining. Three 2×2 or 3×3 200x fields were selected from each mouse and quantified by digital image analysis. Statistical analysis by Student’s T-test comparing mean H-score between vehicle control and high-dose groups for each tissue compartment and each timepoint. (B) Representative photomicrographs of GR (brown) staining counterstained with hematoxylin. Scale bars: 100μm

### Enzalutamide administration reduces the number of granulocytic myeloid-derived suppressor cells within the spleen

Enzalutamide has been previously reported to have potential immunomodulatory effects including increasing thymic weights in a prostate cancer mouse model[23]. Thymic and splenic weights were similar between control and high-dose treated mice at both 8- and 16-weeks (Figure S2A-B). Splenocyte populations in the 8-week treated mice were further evaluated by flow cytometry. The proportion of CD4 and CD8 T-cells were similar in control and high-dose enzalutamide treated mice (Figure S2C). In contrast, there was a reduction in CD11b leukocytes (myeloid-derived suppressor cells [MDSCs]) in mice treated with high-dose enzalutamide relative to control treated mice. Further analysis showed that this reduction of CD11b leukocytes was primarily due to decreased numbers of double-positive Ly6C and Ly6G cells (granulocytic MDSC), while there was no effect on the mononuclear MDSC Ly6C+;Ly6G-population (Figure S2D). These results suggest that enzalutamide may systemically repress the development of granulocytic myeloid-derived suppressor cells.

## Discussion

Enzalutamide is the standard of care for castration-resistant prostate cancer and has also been shown to have therapeutic benefit for castration-sensitive disease[24,25]. Here we used an established endometrial type I cancer mouse model having uterine epithelium specific *Pten* deletion, *Sprr2f-Cre;Pten^fl/fl^*, to evaluate the efficaciousness of enzalutamide in preventing endometrial cancer progression. We demonstrate that enzalutamide, as a single agent, has limited efficacy in decreasing tumor burden in a mouse model of type I endometrial carcinoma. Interestingly, Progesterone Receptor expression in mice treated with enzalutamide was markedly increased, particularly within the stromal compartment. Progesterone antagonizes endometrial epithelial growth through stromal paracrine signaling and stromal Progesterone Receptor expression is critical for this interaction[26]. Thus, we propose that combined treatment with enzalutamide plus progesterone hormone therapy would have synergistic activity in reducing endometrial tumor burden in this mouse model.

Previous xenograft studies testing the efficaciousness of enzalutamide in prostate[14] and breast cancer[27] used castrated and ovariectomized mice, respectively. In these studies investigators observed enzalutamide-dependent decreases in proliferation along with increased apoptosis. However, in our study, the therapeutic effects from enzalutamide treatment were short-lived and restricted to the early induction of apoptosis without any effect on proliferation. These differences in the enzalutamide-dependent regulation of proliferation across individual model systems could be due to complex interactions associated with sex-hormone signaling, influences of microenvironment, or inherent dissimilarities between xenograft studies and genetic models. Proliferation of prostate, breast and endometrial epithelial cells is regulated by sex hormones. Our study mice had an intact reproductive tract and as expected, we observed estrous-dependent effects in proliferation for enzalutamide-treated mice, while the treatment-related effects on apoptosis were not estrous-dependent. It is entirely possible that drug combinations targeting both apoptosis and proliferation might be more efficacious for treating endometrial cancer. Co-administration of progesterone and enzalutamide hormone therapy may produce synergistic effects in decreasing tumor burden by blocking epithelial proliferation through stromal PR signaling and inducing epithelial apoptosis through enzalutamide. Additional studies are warranted to test this combination.

Therapeutic resistance to enzalutamide administration in this model was associated with reduced apoptosis, increased Progesterone Receptor expression, decreased Glucocorticoid Receptor expression and clonal overexpression of p53 protein. Mechanisms for resistance in prostate cancer cells have been associated with impaired apoptosis attributed to increased Glucocorticoid Receptor expression[22]. It is uncertain how this mechanism may apply to our study since GR expression was decreased in resistant cells. Interestingly, two recent clinical studies in prostate cancer have correlated poor treatment response to enzalutamide with p53 alterations in tumors[28,29]. Our data suggest similar ineffectiveness to anti-androgen therapy should be expected in human endometrial cancers containing p53 alterations.

We observed a clear induction of apoptosis and reduction in overall endometrial tumor burden after 8-weeks of treatment. These observations may indicate that AR signaling has an oncogenic role during early stages of endometrial tumorigenesis. Indeed, genetic knockout of murine *Androgen Receptor* combined with heterozygous *Pten* resulted in smaller tumors and fewer number of glands compared to *Pten* knockout alone[30], supporting the oncogenic role of AR in endometrial cancer initiation. This study, however, did not report myoinvasion or metastasis in any of their genetic groups. While initial results measuring tumor load seemed promising, we found that prolonged treatment with enzalutamide did not reduce overall tumor burden and even increased the incidence of progressive disease. We suggest that AR signaling can have both oncogenic and tumor suppressive roles, with tumor initiation being more dependent on oncogenic AR signaling, and later stages of invasion and metastasis dependent on inactivation of tumor suppressive AR signaling.

Studies suggest that enzalutamide may have immunomodulatory effects[23]. We observed no alterations in thymic weights from female mice treated with enzalutamide in contrast to previous observations in male mice treated for prostate cancer[23]. This may represent general differences in immune regulation between males and females. Indeed, gender-dependent biases in immune function and activation have been observed between men and women, including increased immunosuppressive responses in adaptive immunity attributed to testosterone in males[31]. Thus antagonizing AR signaling may produce a more robust immune response in males relative to females. We did observe a decrease in the proportion of splenic granulocytic-MDSCs from treated female mice relative to controls. The functional significance of this finding is uncertain. Increased splenic granulocytic-MDSCs have been associated with cancer in both humans[32] and mice[33]. This observation may reflect, in part, a consequence of sex hormones regulating differentiation of myeloid cells. Indeed, it has been reported that bone marrow cells collected from females have a preference to differentiate along granulocytic lineages, while bone marrow cells from male mice differentiate along monocytic lineages[34]. Determining precisely how modulating AR signaling differentially effects immune function between males and females warrants further investigation, especially since hormone levels frequently change with age and common morbidities like obesity. Thus, the impact of AR regulation on immune function likely has broad implications for a number of human diseases, including cancer, regardless of gender.

In conclusion, we show that enzalutamide administration induces tumor cell apoptosis in a genetic mouse model of endometrial carcinoma. Prolonged administration of enzalutamide results in resistance to apoptosis and clonal expansion of cells expressing stable p53 protein. Our study suggests that the overall efficacy of enzalutamide in treating endometrial carcinoma is restricted to early stages of tumorigenesis.

## Acknowledgements

We thank Jason Bice and Daphne Bryant for assistance with histology. This work was supported by the MUSC HCC Flow Cytometry Shared Resources, the MUSC HCC Biostatistics Shared Resource and the MUSC HCC Translational Sciences Laboratory, supported in part by grant P30 CA138313, National Cancer Institute, Bethesda, MD. This work was funded by National Institutes of Health grants to G.L. (R01CA121275) and a National Comprehensive Cancer Network Oncology Research Program Award to H.M.C. (NCCNENZA0015).

## Declaration of Interests

The National Comprehensive Cancer Network Oncology Research Program and their partners, Medivation and Astellas Pharma, sponsored a portion of this work and reviewed this data, giving permission for its submission for publication.

## Abbreviations

AR: Androgen Receptor
CC3: Cleaved Caspase 3
EMC: Type-I endometrial carcinoma
ER: Estrogen Receptor *alpha*
GR: Glucocorticoid Receptor
MDSC: Myeloid-derived suppressor cell
PR: Progesterone Receptor

---

**Supplemental Figure 1.**
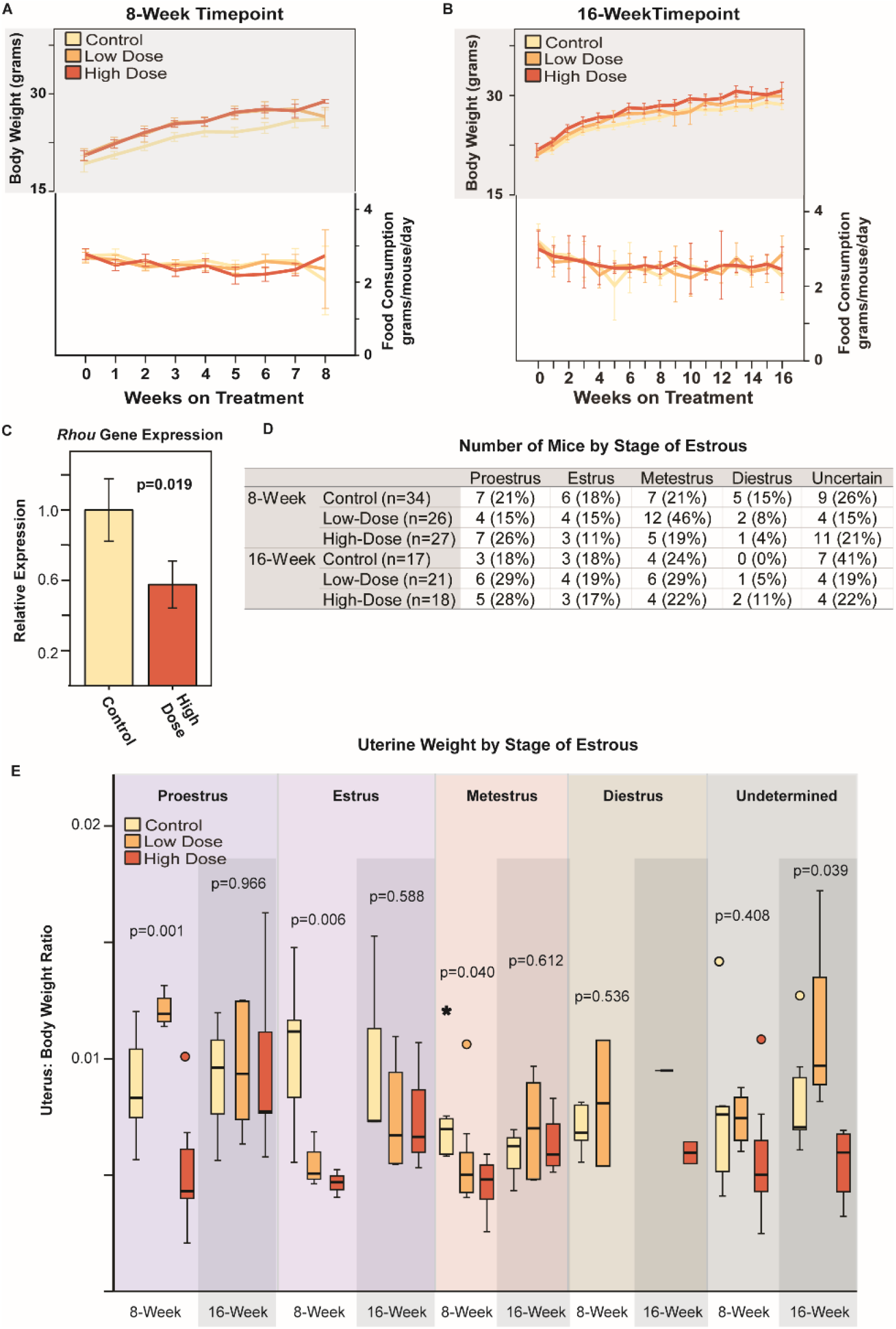
Effects of enzalutamide on body weight, appetite, liver gene expression and uterine weights. (A) 95% confidence intervals for mean body weight (p=0.0332 high-dose vs low-dose, p=0.0046 high-dose vs control and p=0.6262 for low-dose vs control) and daily food consumption (p=0.3626 for high-dose vs low-dose, p=0.4224 for high-dose vs control and p=0.8430 for low-dose vs control) for 8-week timepoint. Statistical analysis by mixed model for repeated measures comparing mean body weight and mean food consumption over time. (B) 95% confidence intervals for mean body weight (p=0.1496 high-dose vs low-dose, p=0.0209 high-dose vs control and p=0.3087 for low-dose vs control) and daily food consumption (p=0.4900 for high-dose vs low-dose, p=0.8811 for high-dose vs control and p=0.5814 for low-dose vs control) for 8-week timepoint. Statistical analysis by mixed model for repeated measures comparing mean body weight and mean food consumption over time. (C) Relative gene expression of AR-target *Rhou* in liver by qPCR (8-week timepoint). Expression levels were normalized to *Rpl19* and calculated by ΔΔCT methods. n=3 mice per treatment group. Statistical analysis by Student’s T-test comparing mean ΔCT for control and high-dose groups. Error bars represent ±2 standard errors of mean. (D) Number of mice for each stage of the estrous cycle based on vaginal histology. (E) Boxplots of uterine:body weight ratios separated by estrous stage. Circles represent outliers that are >1.5 box-lengths beyond the median and asterisks represent outliers that are >3 box-lengths beyond the median. Statistical analysis by one-way ANOVA with post-hoc Bonferroni testing comparing mean ratios for each treatment group and each timepoint.

**Supplemental Figure 2.**
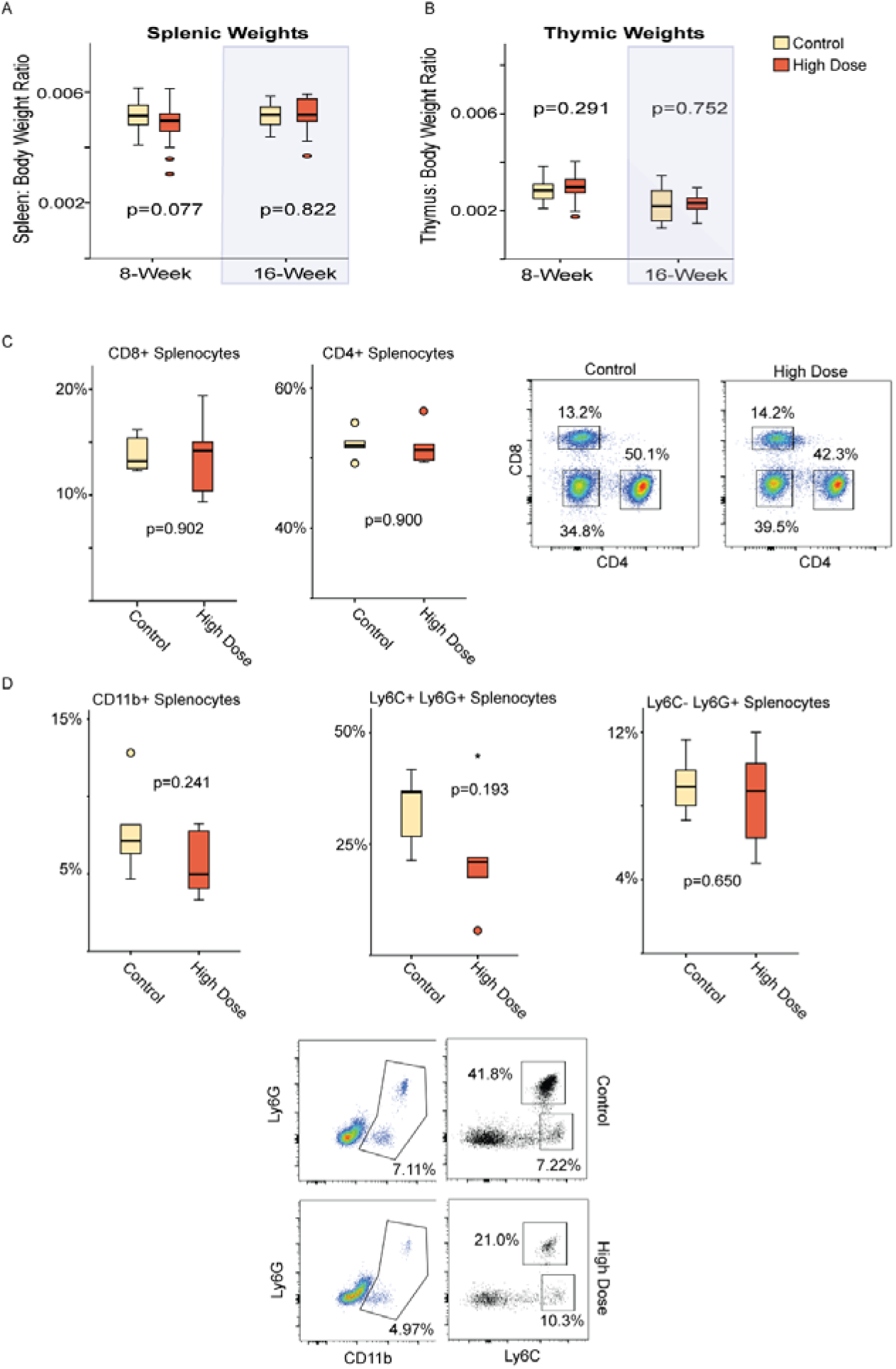
Effects of enzalutamide on spleen and thymus weights, and splenocyte populations. (A-B) Boxplots summarizing individual spleen and thymus weights relative to body weight. (C-D) Flow sorting of CD45+ live splenocytes from study mice (n=5 per dose). Boxplots summarize percentage of given population. Representative gating is depicted in the dotplots. Statistical analysis by Student’s T-test comparing the percentage of positively stained cells between vehicle control and high-dose groups at 8-week timepoint for all the above comparisons.

